# PHYLUCE is a software package for the analysis of conserved genomic loci

**DOI:** 10.1101/027904

**Authors:** Brant C. Faircloth

## Abstract

**Summary:** Targeted enrichment of conserved and ultraconserved genomic elements allows universal collection of phylogenomic data from hundreds of species at multiple time scales (< 5 Ma to > 300 Ma). Prior to downstream inference, data from these types of targeted enrichment studies must undergo pre-processing to assemble contigs from sequence data; identify targeted, enriched loci from the off-target background data; align enriched contigs representing conserved loci to one another; and prepare and manipulate these alignments for subsequent phylogenomic inference. PHYLUCE is an efficient and easy-to-install software package that accomplishes these tasks across hundreds of taxa and thousands of enriched loci.

**Availability and Implementation:** PHYLUCE is written for Python 2.7. PHYLUCE is supported on OSX and Linux (RedHat/CentOS) operating systems. PHYLUCE source code is distributed under a BSD-style license from https://www.github.com/fairclothUlab/phyluce/. PHYLUCE is also available as a package (https://binstar.org/fairclothUlab/phyluce) for the Anaconda Python distribution that installs all dependencies, and users can request a PHYLUCE instance on iPlant Atmosphere (tag: phyluce). The software manual and a tutorial are available from http://phyluce.readthedocs.org/en/latest/ and test data are available from doi: 10.6084/m9.figshare.1284521.

**Supplementary information:** Supplementary Figure 1.

## 1 Introduction

Target enrichment of conserved and ultraconserved elements (hereafter “conserved loci”) allows universal phylogenomic analyses of non-model organisms (Faircloth et al. 2012; Faircloth et al. 2013; Faircloth et al. 2015) at multiple time scales (Faircloth et al. 2012; Smith et al. 2014). The strength of the approach derives from its ability to collect sequence data from thousands of loci across hundreds of species, permitting phylogenetic comparisons across deep phylogenetic breaks such as organismal Classes (> 200-300 Ma) and shallower evolutionary divergences such as populations (< 0.5 – 5 Ma). When the goal of data collection is to infer the evolutionary history of species, the subsequent analytical tasks are generally to: (1) assemble the sequencing reads, which may span tens to hundreds of individuals; (2) identify putative orthologs among the assembled contigs on a sample-by-sample basis while removing putative paralogs; (3) easily generate datasets that contain different individuals, individuals included from other experiments, or individual genome sequences; (4) identify and export sequence data from orthologs across all individuals in the set; (5) align the data and optionally trim resulting alignments in preparation for phylogenetic inference; (6) compute summary statistics on the aligned data; and (7) perform utility functions on the sequence or alignment data prepare them for downstream analyses using a variety of phylogenetic inference programs. PHYLUCE (pronounced “phy-loo-chee”) is the first open-source, easy-to-install software package to perform these tasks for target enriched, conserved loci in a computationally efficient manner.

## 2 Workflow and features

The PHYLUCE workflow (Supplementary Figure 1) for inferring phylogeny begins with external preparation of sequence reads from target-enriched libraries by trimming adapter contamination and low-quality bases using a program like Trimmomatic (Bolger et al. 2014) or a batch processing script similar to illumiprocessor (https://github.com/fairclothlab/illumiprocessor). PHYLUCE then offers several programs to batch-assemble the resulting “clean” reads into contigs using different assembly programs (Zerbino and Birney 2008; Simpson et al. 2009; Grabherr et al. 2011) with parallelization approaches tailored to each program. The next step in the PHYLUCE workflow is to identify orthologous conserved loci shared among individuals. The match_contigs_to_probes program performs the steps of ortholog identification and paralog removal by aligning the assembled contigs to a FASTA file of target enrichment baits using lastz (Harris 2007). Although this program is designed to work with standardized baits sets developed for the targeted enrichment of UCE loci (e.g. http://ultraconserved.org), users can input custom bait sets with different naming conventions targeting different classes of loci by adjusting several parameters (e.g., Mandel et al. 2014). Following the alignment step, match_contigs_to_probes screens the lastz output to identify (1) assembled contigs hit by probes targeting different loci, and (2) different contigs that are hit by probes targeting the same locus. The program assumes that these reciprocally duplicate loci are potentially paralagous and removes them from downstream analytical steps. The program then builds a relational database containing a table of detections and non-detections at each locus across all input assemblies as well as a table associating the name of each targeted locus (from the FASTA file representing the bait set) with the name of the assembled contig to which it matches. Next, users of PHYLUCE create a “taxon-set” configuration file that specifies the individual assemblies that will be used in downstream phylogenetic analyses. By inputting this configuration file to the get_match_counts program, users can flexibly create different data sets, integrate data from separate studies targeting the same loci, or include identical loci harvested from published genome sequences (e.g. http://github.com/faircloth-lab/uce-probe-sets). After identifying those individuals and loci in the desired taxon set, users extract the contigs corresponding to no^duplicate conserved loci into a monolithic (all loci for all taxa) FASTA-formatted file using the get_fastas_from_match_counts program. This program renames each contig for each species within the taxon set such that the FASTA header for each contig contains information denoting the species in which the conserved locus was detected and the specific conserved locus to which it matched. After creating the monolithic FASTA, users can align the targeted loci with the seqcap_align program, which parallelizes MAFFT (Katoh and Standley 2013) or MUSCLE (Edgar 2004) alignments across all targeted loci on computers with multiple CPUs. The seqcap_align program also offers the option to trim the resulting alignments for edges that are poorly aligned - a suitable choice when the species within the taxon set are closely related (e.g., roughly Orde-level or lower taxonomic ranks, < 50 Ma). To apply more aggressive alignment trimming when relationships are older (> 50 Ma), PHYLUCE provides a similar program that implements parallelized, internal trimming using Gblocks (Castresana 2000; Talavera and Castresana 2007).

PHYLUCE includes several parallelized programs to manipulate the resulting alignments, including the ability to rapidly generate summary statistics across thousands of alignments, explode alignments into their corresponding FASTA sequences, extract taxa from alignments, compute parsimony informative sites within alignments, and convert alignments between common formats, and these programs can also be used with alignments from other data types. After alignment, PHYLUCE users can generate data matrices having varying levels of completeness using the get_only_loci_with_min_taxa program. This program screens each locus for taxonomic completeness and filters out loci containing fewer taxa than desired. In this way, users can create 100% complete (all taxa have data for all loci) or incomplete data matrices (some loci have data for a certain percentage of taxa). After filtering loci for taxonomic completeness, PHYLUCE offers several programs to format resulting alignments for analyses in PartitionFinder (Lanfear et al. 2012), RAxML (Stamatakis 2014), ExaBayes (Aberer et al. 2014), GARLI (Zwickl 2006), or MrBayes (Ronquist and Huelsenbeck 2003). Programs are also available to assist users with preparing data for and running gene-tree-based species tree analyses.

## Acknowledgements

I thank Carl Oliveros, Nick Crawford, and Mike Harvey for contributing to the source code and Travis Glenn, John McCormack, Michael Alfaro, Robb Brumfield, Brian Smith, and Kevin Winker for contributing to early UCE studies. Comments from David Posada and three anonymous reviewers improved this manuscript.

---

**Supplementary Figure 1.**
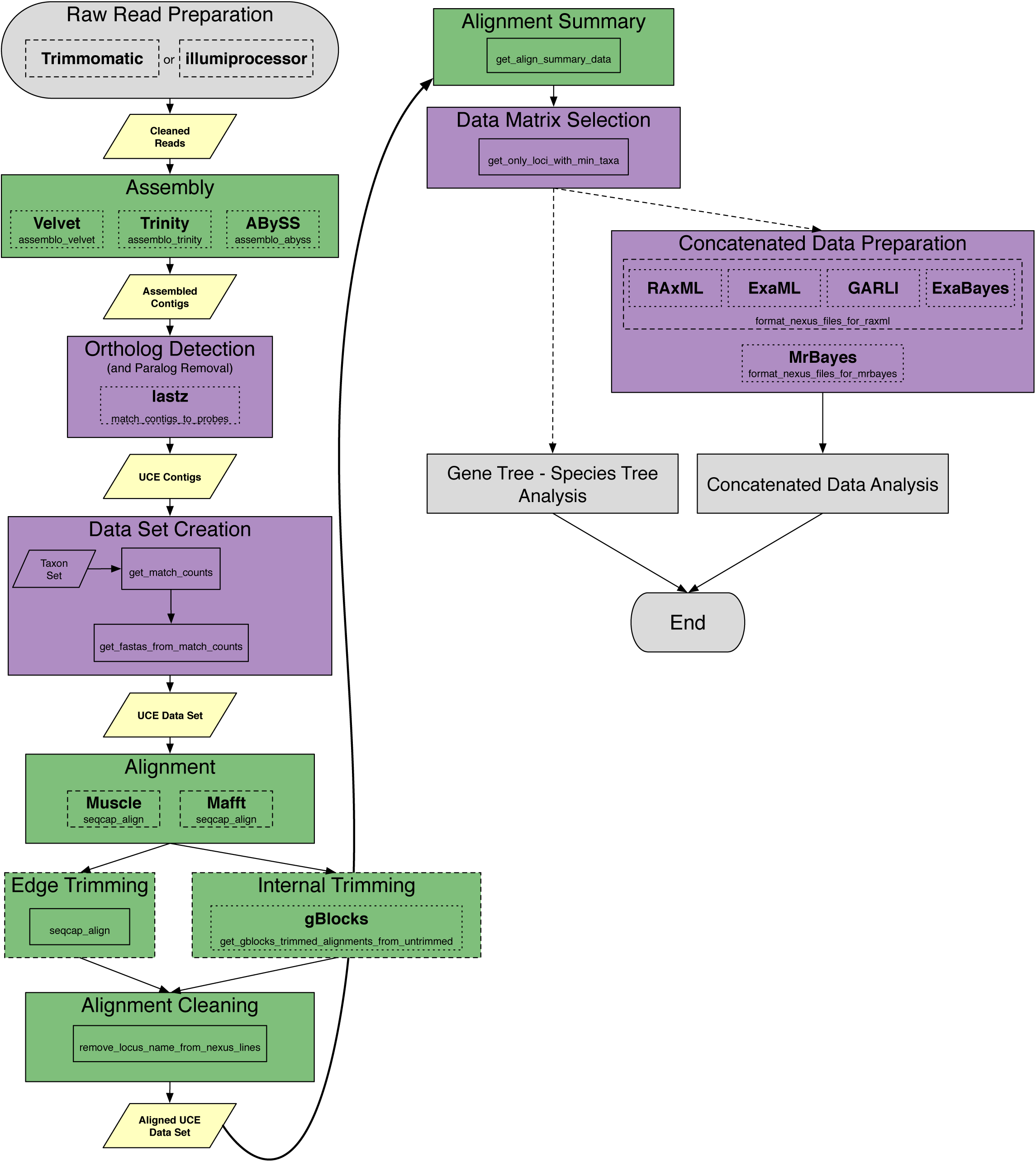
PHYLUCE workflow for phylogenomic analyses of data collected from conserved genomic loci using targeted enrichment.

